# Multiplexed CRISPR-Cas9 based genome editing of *Rhodosporidium toruloides*

**DOI:** 10.1101/545426

**Authors:** Peter B. Otoupal, Masakazu Ito, Adam P. Arkin, Jon K. Magnuson, John M. Gladden, Jeffrey M. Skerkerd

**Affiliations:** Joint BioEnergy Institute, Emeryville, California, USA; Biomass Science and Conversion Technologies, Sandia National Laboratories, Livermore, California, USA; T-Frontier Division, Frontier Research Center, Toyota Motor Corporation, Aichi, Japan; Energy Biosciences Institute, Berkeley, California, USA; Environmental Genomics and Systems Biology Division, Lawrence Berkeley National Laboratory, Berkeley, California, USA; Biological Systems and Engineering Division, Lawrence Berkeley National Laboratory, Berkeley, California, USA; Department of Bioengineering, University of California, Berkeley, Berkeley, California, USA; Chemical and Biological Processing Group, Pacific Northwest National Laboratory, Richland, Washington, USA

**Author notes:** Address correspondence to Peter B. Otoupal, or Jeffrey M. Skerker.

**Keywords:** *Rhodosporidium toruloides*, CRISPR-Cas9, genome engineering, multiplexed, *URA3*, *CAR2*, tRNA

## Abstract

Microbial production of biofuels and bioproducts offers a sustainable and economic alternative to petroleum-based fuels and chemicals. The basidiomycete yeast *Rhodosporidium toruloides* is a promising platform organism for generating bioproducts due to its ability to consume a broad spectrum of carbon sources (including those derived from lignocellulosic biomass) and to naturally accumulate high levels of lipids and carotenoids, two biosynthetic pathways that can be leveraged to produce a wide range of bioproducts. While *R. toruloides* has great potential, it has a more limited set of tools for genetic engineering relative to more advanced yeast platform organisms such as *Yarrowia lipolytica* and *Saccharomyces cerevisiae.* Significant advancements in the past few years have bolstered *R. toruloides’* engineering capacity. Here we expand this capacity by demonstrating the first use of CRISPR-Cas9 based gene disruption in *R. toruloides.* Stably integrating a Cas9 expression cassette into the genome brought about successful targeted disruption of the native *URA3* gene. While editing efficiencies were initially low (0.002%), optimization of the cassette increased efficiencies 364-fold (to 0.6%). Applying these optimized design conditions enabled disruption of another native gene involved in carotenoid biosynthesis, *CAR2,* with much greater success; editing efficiencies of *CAR2* deletion reached roughly 50%. Finally, we demonstrated efficient multiplexed genome editing by disrupting both *CAR2* and *URA3* in a single transformation. Together, our results provide a framework for applying CRISPR-Cas9 to *R. toruloides* that will facilitate rapid and high throughput genome engineering in this industrially relevant organism.

**IMPORTANCE:** Microbial biofuel and bioproduct platforms provide access to clean and renewable carbon sources that are more sustainable and environmentally friendly than petroleum-based carbon sources. Furthermore, they can serve as useful conduits for the synthesis of advanced molecules that are difficult to produce through strictly chemical means. *R. toruloides* has emerged as a promising potential host for converting renewable lignocellulosic material into valuable fuels and chemicals. However, engineering efforts to improve the yeast’s production capabilities have been impeded by a lack of advanced tools for genome engineering. While this is rapidly changing, one key tool remains unexplored in *R. toruloides*; CRISPR-Cas9. The results outlined here demonstrate for the first time how effective multiplexed CRISPR-Cas9 gene disruption provides a framework for other researchers to utilize this revolutionary genome-editing tool effectively in *R. toruloides.*

---

*Rhodosporidium toruloides* (also known as *Rhodotorula toruloides)* is a basidiomycetous yeast that has attracted interest for its great bioengineering potential. The oleaginous yeast can be readily cultivated in high-density cell cultures (1), and naturally accumulates large quantities of both carotenoids and lipids (2) which serve as precursors for many valuable compounds. Complementing this is the ability of *R. toruloides* to efficiently consume a wide variety of carbon sources, including those found in lignocellulose hydrolysates (3, 4). Furthermore, *R. toruloides* can grow robustly under difficult environmental conditions including high osmotic stress (5) and the presence of ionic liquids used in pretreatment processes (6–8). It is therefore being assessed as a novel platform for the industrial-scale generation of valuable commodities including biofuels, pharmaceuticals, and other advanced bioproducts (9–12).

Despite these advantages, commercial adoption of *R. toruloides* will be hampered until effective genetic engineering tools are developed in this system (13–15). Notably, no autonomously replicating sequences (ARS) have been identified for use in *R. toruloides,* meaning that all genetic manipulation of this organism to date has been accomplished by direct integration of heterologous DNA elements into the genome. The genetic engineering toolkit for *R. toruloides* has been expanded in recent years with the development of transformation techniques such as *Agrobacterium tumefaciens-*mediated transformation (16) (ATMT), electroporation (15, 17) and lithium acetate chemical transformation (14), which have enabled both efficient random integration based on the non-homologous end joining (NHEJ) pathway, and targeted deletion/integration based on homologous recombination (HR) (18, 19),

Concurrent with these advances is a revolution in the field of genome engineering brought on by the development of precise genome and transcriptome editing through clustered regularly interspaced short palindromic repeat (CRISPR)-based systems (20–22). Modern genetic engineering approaches in other organisms have largely gravitated towards using CRISPR systems to enact desired DNA changes (23–25). This approach utilizes a ribonucleoprotein complex (RNP) consisting of a CRISPR associated nuclease (Cas, the most common being Cas9) and a synthetic single guide RNA (sgRNA). The sgRNA contains a ∼20 nucleotide (nt) spacer sequence complementary to a unique DNA sequence in the targeted organism, as well as a 76 nt handle that complexes with Cas9. By modifying the 20 nt spacer sequence of the sgRNA, researchers can direct the nuclease to a specific location and enact targeted DNA cleavage.

While CRISPR-Cas9 has revolutionized the world of genome editing, it has yet to be demonstrated in *R. toruloides.* The yeast’s potential as a robust host for the production of bioproducts would be significantly improved by the development of CRISPR-Cas9 strategies for engineering its genome, especially considering that the genetic engineering toolkit for *R. toruloides* has fallen behind that of more developed yeasts such as *Yarrowia lipolytica* (26–29). Cas9 engineering is more amenable to multiplexed gene editing than the current techniques for manipulating *R. toruloides’* genome based on ATMT (30). Furthermore, CRISPR-Cas edits can be employed in a matter of days (31), while ATMT gene editing can take anywhere from two weeks to one month (16). Another advantage of CRISPR-Cas9 gene editing is that it can utilize NHEJ to create site-specific gene deletions, while current techniques relying on HR via *KU70* deletion of the NHEJ repair pathway suffer from the apparent low activity of HR repair relative to NHEJ in *R. toruloides* (15, 32). Targeted gene disruptions would likely be more easily accomplished utilizing CRISPR-Cas9 targeting followed by error-prone NHEJ repair than the current approach.

Here we demonstrate the first application of gene editing using CRISPR-Cas9 in *R. toruloides.* Genomic integration of the coding sequences of Cas9 and an associated sgRNA allows for targeted gene disruption of *URA3* upon (likely NHEJ-based) repair of DNA cleavage. Although initial editing efficiency was low (0.0017% ± 0.0011%), optimizing expression of the sgRNA and fine-tuning of its target sequence led to a 364-fold improvement in editing success of up to 0.62% ± 0.50%. Using the design principles learned in targeting *URA3,* we developed CRISPR-Cas9 constructs to disrupt the carotenoid biosynthesis gene, *CAR2.* Attempts to delete this gene resulted in significantly greater success, with editing efficiencies reaching up to 46 ± 22% of all cells transformed with the CRISPR-Cas9 expression cassette. We further show that multiplexed gene disruption of the *R. toruloides* genome is possible using this approach. Combining an array of four sgRNAs separated by self-processing tRNA elements successfully allowed for targeted deletions of both *CAR2* and *URA3* in one simultaneous transformation. We observed both indels near each cut site and complete deletion of the region between the cut sites. We demonstrate that by using two sgRNAs to disrupt each gene, the intergenic region between these two cut sites can be excised during the DNA repair process. Interestingly, *R. toruloides* demonstrated a propensity to re-insert the excised DNA in its reverse orientation, further supporting the observation that the organism is particularly effective at accomplishing NHEJ-based DNA repair. Taken together, these results outline a strategy for achieving efficient CRISPR-based genome editing in *R. toruloides* and will streamline metabolic engineering efforts in this industrially relevant organism.

## RESULTS

### Integration of Cas9 and sgRNA into *R. toruloides’* genome allows for targeted gene disruption

We first designed a Cas9 expression construct for use in *R. toruloides.* This included codon optimization, addition of a nuclear localization sequence (NLS) to Cas9, and selection of the native GAPDH promoter for constitutive expression. An sgRNA expression cassette was placed upstream of the Cas9 sequence cassette consisting of a promoter, a hepatitis delta virus (HDV) ribozyme cleavage sequence, a 20 nt unique gene targeting sequence, a 76 nt common sgRNA handle for the association of Cas9 with the RNA, and a terminator sequence. The SNR52 RNA polymerase III promoter and terminator sequences from *Saccharomyces cerevisiae* were employed to drive expression of this sgRNA, as they have been used successfully in other fungi (33, 34). Placing a ribozyme element between the SNR52 promoter and the sgRNA element resulted in improved editing efficiency in S. *cerevisiae* in our previous work (35). The ribozyme was therefore included in the hopes of increasing editing efficiency in *R. toruloides.*

To validate this CRISPR-Cas9 system, the *URA3* gene encoding orotidine 5’-phosphate decarboxylase was targeted for deletion. Expression of the decarboxylase is known to allow yeast to convert 5-fluoroorotic acid (5-FOA) into 5-fluorouracil, a compound that is highly toxic to most yeast (36). Therefore, successful editing (i.e. loss of function of *URA3* caused by error-prone NHEJ repair of the Cas9-mediated dsDNA break) can be selected for by growth in the presence of 5-FOA.

The first attempt to generate edits at the *URA3* locus was done in a single-step by transforming a PCR fragment containing both the Cas9 and a *URA3-*specific sgRNA expression cassette, followed by selection on 5-FOA plates (Fig. 1A). The number of 5-FOA resistant (*5-FOA*^*R*^) colonies obtained in the presence of the Cas9/sgRNA cassette was indistinguishable from a control transformation in the absence of the PCR fragment, suggesting that editing, if it occurred, was no more frequent than the rate of spontaneous 5-FOA^R^ (Fig. 1B). Sequencing DNA from three 5-FOA^*R*^ colonies near the cut site of Cas9 revealed a consistent frame shift occurred outside of the target sequence (Fig. 1C). This suggests that *5-FOA*^*R*^ arose not from Cas9-mediated gene disruption, but through spontaneous mutation of *URA3* leading to loss of function. We thought that this failure in Cas9-based gene editing may have been due to poor expression of the sgRNA from the non-native SNR52 promoter sequence, or due to targeting an area of the genome not amenable to Cas9 binding. However, even upon replacing the SNR52 promoter with four other variants, and utilizing an alternative sgRNA, no improvement in 5-FOA colony formation was observed (Fig. S1). Indeed, each of these variants resulted in the same spontaneous mutation outside of the Cas9 cut site (Fig. S1).

**FIG 1.**
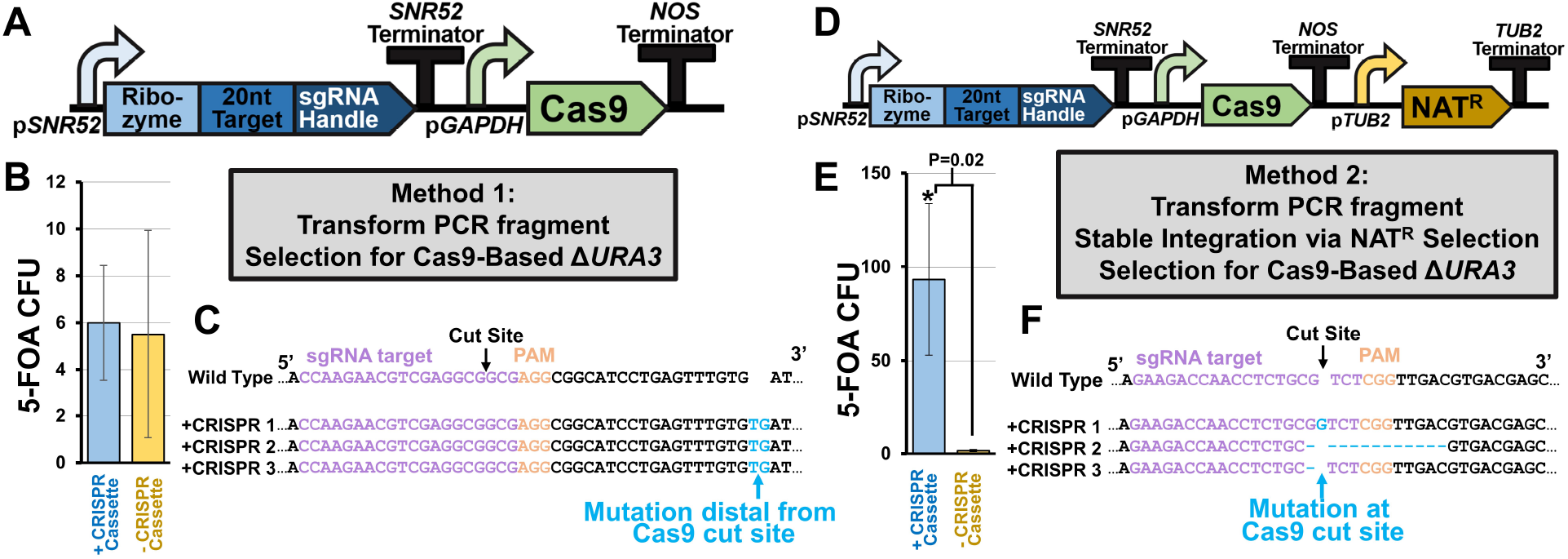
Targeted gene disruption using CRISPR-Cas9. (A) Schematic of original CRISPR-Cas9 design for causing indels in *R. toruloides.* A PCR fragment containing the coding sequences for expressing sgRNA and Cas9 is transformed into competent cells, which uptake the DNA into their nucleus and express the machinery from the PCR fragment. (B) Total colony forming units (CFU) of *R. toruloides* under 5-FOA selection with and without application of this CRISPR-Cas9 editing scheme. (C) Partial sequencing of *URA3* of each potentially edited colony near the cut site of Cas9. (D) Revised protocol in which the coding sequence for a selectable marker is included in the PCR fragment and an additional selection step for integration of the fragment into the genome is included. (E) Total CFU of *R. toruloides* under 5-FOA selection with this revised CRISPR-Cas9 editing scheme. Significance was calculated using a two-tailed type II student’s *t*-test. (F) Partial sequencing of *URA3* of three edited colonies near the cut site of Cas9. All error bars represent the standard deviation of biological triplicates.

It was hypothesized that this failure of Cas9-mediated gene editing was due to insufficient expression of the CRISPR machinery during the transformation process. To alleviate this problem, a new approach was employed in which the CRISPR cassette was first targeted for stable integration using a selectable drug marker cassette that encodes for nourseothricin resistance *(NAT*^*R*^) before screening for successful *URA3* editing (Fig. 1D).

Approximately 300 *NAT*^*R*^ colonies were obtained after transformation and three colonies were selected for subsequent screening of Cas9-mediated DNA editing. Growing each colony in YPD followed by plating on YPD supplemented with 5-FOA resulted in 93 ± 40 colonies (Fig. 1E). A control transformation of *NAT*^*R*^ without the CRISPR cassette resulted in significantly fewer spontaneously 5-FOA^R^ colonies (1.6 ± 0.5) This was significantly above the background level of spontaneous 5-FOA resistance observed utilizing the same process in the control transformation with no PCR DNA (P = 0.02). Furthermore, sequencing revealed indels near the cut site that resulted in frameshifts, suggesting that error-prone NHEJ repair occurred in a specific location dictated by the sgRNA sequence (Fig. 1F). Utilizing the other four promoter variants also resulted in successful gene disruption at the Cas9 cut site (Fig. S2).

These results indicate that stable integration of a Cas9-sgRNA expression cassette into the genome allows for successful gene editing in *R. toruloides.* Additionally, growth curves of wild-type cells and cells harboring the Cas9-sgRNA cassette show no difference in growth rates, indicating that the cassette does not elicit detrimental fitness effects (Fig. S3). However, while the CRISPR-Cas9 system was able to disrupt genes, the efficiency of this process was low. The efficiency of gene editing was determined by comparing the total amount of colony forming units (CFUs) in the presence of 5-FOA to CFUs in the absence of 5-FOA. Total colony forming units were roughly 10^5^-fold higher in the absence of 5-FOA, indicating that Cas9 editing efficiency was on the order of ∼0.001%. Due to this overall low editing efficiency, focus shifted to the optimization of editing efficiency. Initial tests attempting to improve Cas9-editing by adding an additional NLS resulted in a decrease in editing efficiency (Fig. S4), so improvement in the design of the sgRNA sequence was pursued in the following experiments.

### sgRNA target optimization

The first point of optimization in the sgRNA design was to re-consider the 20nt DNA targeting sequence used in the sgRNA. Complex secondary structure near the DNA target sequence can lead to significant hindrance to Cas9 activity (37). Therefore, the program sgRNA Scorer 2.0 was used to optimize target sequences of Cas9 (38). This program is based on sgRNA design principles uncovered in human cell lines, and while it has been utilized to design successful sgRNAs for Cas9 editing in yeast (39), its applicability outside of human cell lines is not well-known. Therefore, seven new target sequences for *URA3* were selected representing a range of different predicted scores and a series of new editing constructs were generated (plasmids 213-233, Table S1) to test their relative editing efficiency. Altering the sgRNA targeting sequence had a noticeable impact on editing efficiency. In the optimal case, editing efficiency was improved ∼14-fold over the original sgRNA target sequence, while the sgRNA predicted to perform worst reduced editing efficiency ∼9-fold (Fig. 2A). This indicates that sgRNA target optimization is important for achieving acceptable levels of editing efficiency in *R. toruloides,* and that established design tools can facilitate construction of new sgRNA targets.

**FIG 2.**
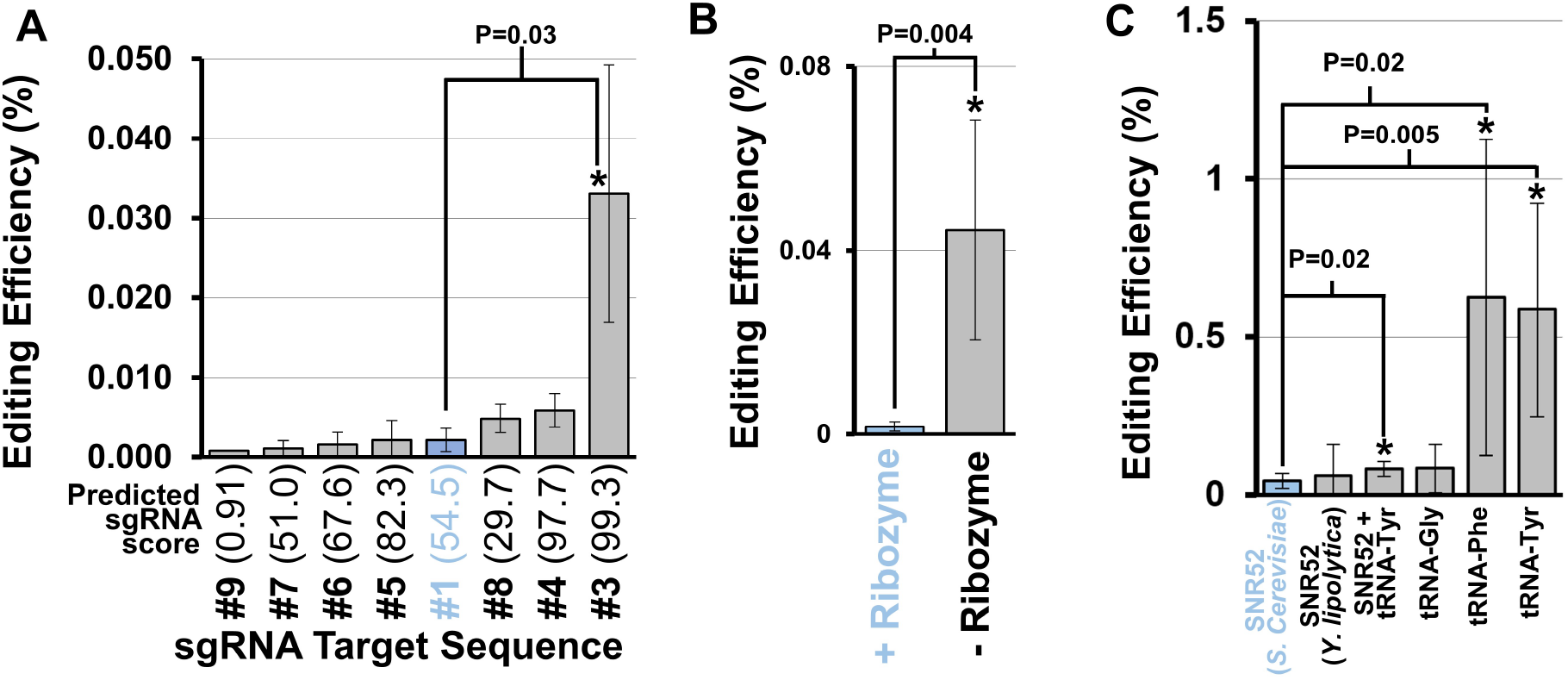
Optimization of sgRNA expression and target sequence. (A) Editing efficiency of various sgRNA target sequences. Bars indicate measured CRISPR-Cas9 gene editing efficiency, as a percentage of the total cells in the transformation mixture exhibiting the expected edited phenotype. Significant differences from the original target sequence (highlighted in blue, P < 0.05) is indicated by asterisks and calculated using a two-tailed type II student’s *t*-test. The predicted aptitude for a target sequence to achieve successful DNA editing based on the sgRNA Scorer 2.0 algorithm (38) is depicted in parentheses after each target sequence. (B) Measured editing efficiency of sgRNA with and without an HDV ribozyme cleavage site included between the promoter and the 20 nt target sequence. (C) Measured editing efficiency of various promoters used to drive sgRNA expression. All asterisks indicate statistical difference from the original expression design (highlighted in blue, P < 0.05) calculated using a two-tailed type II student’s *t*-test. Error bars represent the standard deviation of biological triplicates.

Another point of optimization was the ribozyme included between the sgRNA promoter and the 20nt guide sequence. The ribozyme was originally included for its potential to protect the 5’ end of the sgRNA from 5’ exonucleases, and was found to aid in improving editing efficiencies in S. *cerevisiae* (35, 40). The ribozyme was removed to see if this was the case. This alteration caused the editing efficiency to increase 26-fold (Fig. 2B), significantly improving Cas9-directed gene editing (P = 0.004). As such, we recommend the exclusion of the 5’ HDV ribozyme in designing sgRNAs for expression in *R. toruloides.*

Alternatives to the promoter sequence used in driving sgRNA expression were explored next. A variety of RNA Pol-III promoters were examined, each excluding the detrimental ribozyme element. The original SNR52 promoter used here, which was originally derived from *S*. *cerevisiae*, was replaced with an analogous SNR52 promoter element from another oleaginous yeast, *Y. lipolytica,* as this sequence has been proven to produce functional sgRNAs in other systems (41). However, this change made no significant impact on editing efficiency (Fig. 2C). Since it is unknown if SNR52 exists in *R. toruloides*, a native SNR52 sequence for driving sgRNA expression could not be used.

To utilize a native *R. toruloides* sequence, we turned to an alternative promoter system to drive sgRNA expression. Work in other fungi has found that tRNAs can serve as promoters for sgRNAs *in vivo* (35, 42). Furthermore, tRNAs contain internal elements that promote RNase P and Z mediated cleavage at specific sites, allowing for formation of precise final sgRNA sequences. Including the *R. toruloides* tRNA^Tyr^ sequence downstream of the S. *cerevisiae* SNR52 promoter increased editing efficiency slightly (1.8-fold), but significantly (P = 0.02) (Fig. 2C). Furthermore, directly replacing the SNR52 promoter with tRNAs led to successful editing. This was particularly true of tRNA^Phe^ and tRNA^Tyr^, whose use as sgRNA promoters led to editing efficiencies 14-fold and 13-fold greater, respectively (or 0.62% ± 0.50% and 0.59% ± 0.34% editing efficiency, respectively). These results indicate that native *R. toruloides* tRNA promoters are more effective than heterologous SNR52 promoters for Cas9-based gene editing. This could be due to increased sgRNA expression, although this remains to be tested.

### Multiplexed gene disruption with CRISPR Cas9

In order to develop a multiplex gene editing system for *R. toruloides, a* second target gene was selected, *CAR2,* a gene that encodes for a phytoene synthase/lycopene cyclase protein that is essential for carotenoid biosynthesis (43). Loss of *CAR2* function is easily observed as a change in colony color from red to white.

A set of single guides targeting *CAR2* were first designed to disrupt the *CAR2* locus and their editing efficiencies tested. Four different Cas9-sgRNA-*NAT*^*R*^ constructs were built using the design principles discovered for *URA3* editing and transformed into *R. toruloides.* After stable integration using the *NAT*^*R*^ selection method and replating, a significant number of white colonies for all four sgRNA variants were observed. Editing efficiencies (determined as the ratio of white to red colonies) ranged from 3.4 ± 2.7% to 46.2 ± 22.2% (Fig. 3A). Notably, these levels of editing efficiency are substantially higher for *CAR2* than for any of the *URA3* targeting constructs, indicating that this genome region is more amenable to Cas9-based genome editing. Furthermore, successful disruption of *CAR2* indicates that multiplexed gene editing might be possible by selecting for *5-FOA*^*R*^ colonies that exhibit a white phenotype.

**FIG 3.**
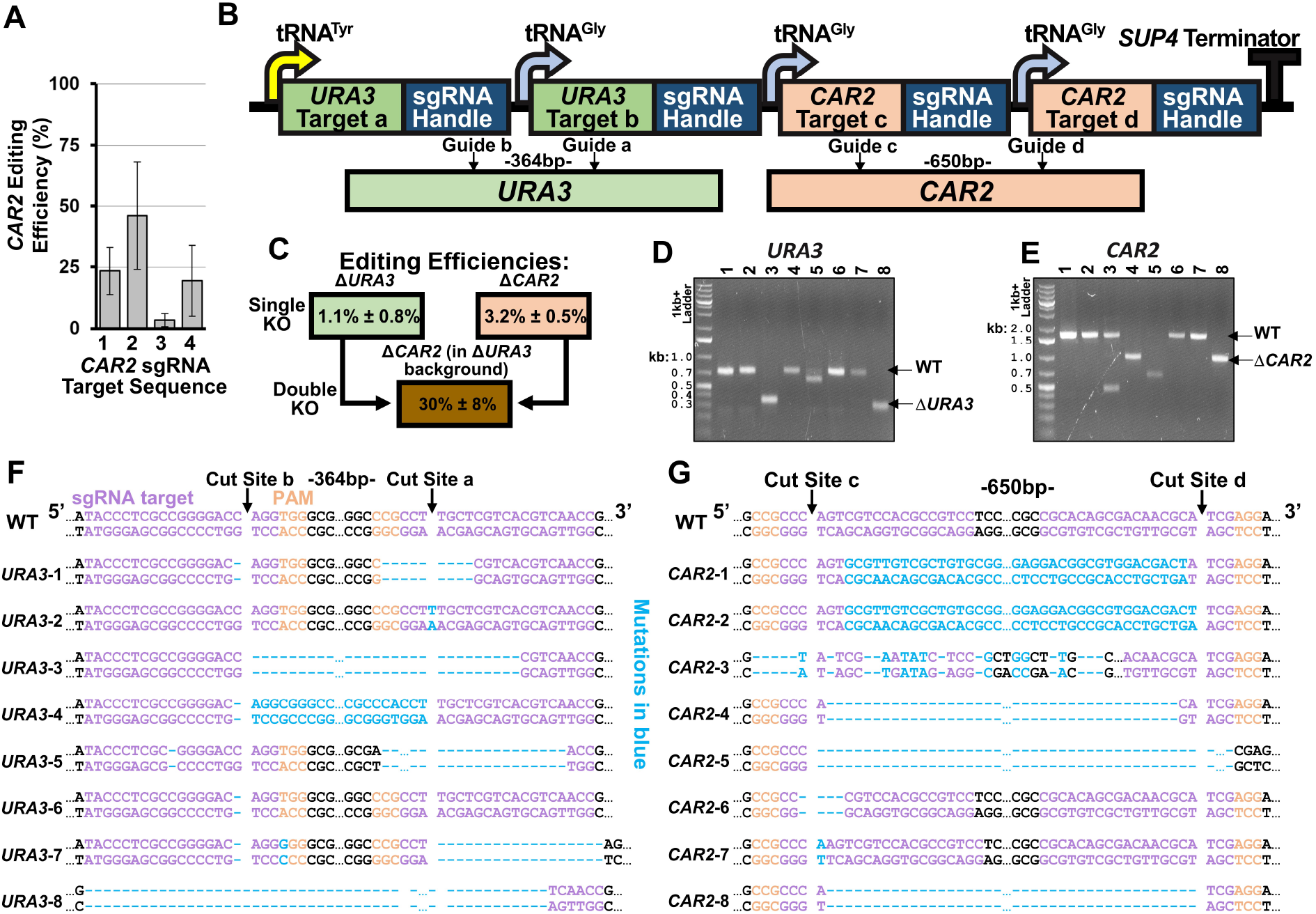
Multiplexed gene disruption using CRISPR-Cas9 in *R. toruloides.* (A) Editing efficiency of four sgRNAs targeting *CAR2.* Error bars represent standard deviations of biological triplicates. (B) The design used to express multiple sgRNAs in a single array. The specific cut sites for *URA3* and *CAR2* are shown below. Guides “a” and “b” correspond to sgRNAs “4” and “3” for *URA3,* while guides “c” and “d” correspond to sgRNAs “1” and “3” for *CAR2* based on the sequences presented in Table S3. (C) 5-FOA plate with colonies demonstrating successful simultaneous disruption of both genes. To the right is shown the editing efficiency of disrupting each gene individually, as well as tandem gene-disruption editing efficiency. (D, E) Gel image showing PCR amplification of genomic DNA of eight unique colonies near the targeted cut site of (D) *URA3* or (E) *CAR2.* (F) Sequencing results near the target cut site of the eight *URA3* PCR products from (D). Cas9 cut sites are indicated, as well as the DNA size of the excised DNA fragment between each cut site. Mutations are highlighted in blue. (G) Sequencing results near the target cut site of the eight *CAR2* PCR products from (E). Mutations caused by Cas9 targeting are highlighted in blue. All error bars represent standard deviation of biological triplicates.

To explore multiplexed deletion of two genes in *R. toruloides,* a Cas9 construct targeting both *URA3* and *CAR2* was created. For this, multiple sgRNAs were placed together sequentially into an array, with each guide RNA separated by a tRNA sequence (Fig. 3B). This approach has been applied in other organisms to take advantage of inherent tRNA post-transcriptional processing to express multiple unique sgRNA sequences (42, 44, 45). Combining multiple sgRNAs in an array is particularly useful for *R. toruloides,* as this minimizes the amount of genetic material that needs to be delivered while maximizing the number of potential gene targets. Additionally, utilizing multiple sgRNAs to target one gene at different locations allows for the possibility of removing a large DNA in between the target sites, and also increases the possibility that the target gene is successfully disrupted (20, 46, 47). Therefore, our constructs were designed to express four sgRNAs (two for each gene) such that cleavage would occur at two sites separated by ∼500bp in both genes.

The multiplexed CRISPR editing construct fragment was transformed into *R. toruloides* and multiple NAT^R^ colonies were selected to screen for genetic disruptions. To determine loss of *CAR2* function, we grew colonies overnight in liquid cultures, which were subsequently plated on Nat plates. Colonies were then screened for their red or white phenotypes. This provided an estimate of the editing efficiency of *CAR2,* which was determined to be 3.2% ± 0.5%, independently of the editing efficiency of *URA3.* To determine loss of *URA3* function, an equal volume of each culture was plated on both Nat and 5-FOA plates and the CFU counts on each plate were compared. This provided an estimate of the editing efficiency of *URA3* of 1.1% ± 0.8%. Colonies growing on 5-FOA were also screened for their red or white phenotypes to determine the dual-gene disruption efficiency. Of the *5-FOA*^*R*^ colonies, 30.0% ±8.0% exhibited a white phenotype, indicating simultaneous *CAR2* and *URA3* disruption. A representative example of a 5-FOA plate demonstrates the screening of dual-gene disruption (Fig. 3C).

We next sought to confirm that these edits were indeed the result of successful Cas9-mediated editing. For this, eight white colonies were selected from the 5-FOA plates and the regions of *URA3* and *CAR2* which surrounded the Cas9 target sites were PCR amplified. Gel electrophoresis of these PCR products revealed that several of the samples reduced significantly in size from the predicted size of the wildtype PCR products (717nt and 1637nt for *URA3* and *CAR2* respectively) (Fig. 3 D, E).

Sequence-verification of these fragments was employed around the target cut sites to see what type of gene editing events occurred (Fig. 3 F, G). Sequencing revealed that most of the cut sites contained indels resulting in frameshift mutations. Cas9-based gene editing was observed at both target sites for each gene in seven out of eight replicates. Replicates three and eight demonstrated complete excision of the intergenic region between both cut sites, while replicate four surprisingly demonstrated re-integration of the intergenic region in its reverse direction. Similarly, editing of *CAR2* occurred in every sample at cut site three, and at cut site four in all samples excluding replicates three, six, and seven. Replicates four, five, and eight demonstrated complete removal of the intergenic region between both cut sites. Again, replicates one and two demonstrated the surprising evidence of re-integration of the reverse direction of the intergenic region. Taken together, these results demonstrate that multiplexed gene disruption mediated by CRISPR-Cas9 is possible in a single transformation of *R. toruloides.*

## DISCUSSION

The development of CRISPR-Cas9 technology has revolutionized genome engineering. Scientists are now able to rapidly edit the DNA of organisms where genetic manipulation was previously inefficient or intractable (48). The fundamental science underlying CRISPR engineering of genomes remains constant regardless of the organism being investigated; scientists use these nucleases to induce DNA cuts at highly specific locations and rely upon the organism’s DNA repair machinery to mend these breaks with donor DNA (via HR) or in an error-prone fashion (via NHEJ). However, the process of delivering and expressing fully functional CRISPR components, and the biology of each organism’s DNA repair pathways, requires organism-specific optimization to accommodate each species’ unique characteristics. The past two years have seen the publication of a wide array of studies demonstrating CRISPR-Cas9 genome engineering in fungal species for which few robust DNA editing tools existed, including *Aspergillus niger* (49), *Cryptococcus neoformans* (50), *Mucor circinelloides* (51), and *Myceliophthora thermophila* (47). The dawning of a “fungal CRISPR revolution” has occurred, empowering researchers to explore new bioproduction possibilities in obscure yet promising fungi.

Here we add *R. toruloides* to the list of fungi now editable using CRISPR-Cas9. While this yeast has been touted for its great bioproduction potential, the sparse genetic manipulation toolkit relative to other organisms such as *Y. lipolytica* (27) previously hindered engineering efforts (52–55). The past four years have seen various researchers remedy this problem with the development of tools for transforming *R. toruloides* and efficiently expressing exogenous DNA (15, 56). While the toolkit has expanded to include useful promoters (53), drug markers (11), and targeted gene editing methods (18), CRISPR-Cas9 methods for advanced genome engineering have been lacking. This study outlines the strategies by which researchers can employ multiplexed CRISPR-Cas9 genome editing to manipulate *R. toruloides.* These are the first steps to ultimately achieving more sophisticated genome- or transcriptome-scale engineering of *R. toruloides,* an important step towards fulfilling the organism’s potential.

Accomplishing this required overcoming significant barriers. Most notably, the lack of a plasmid capable of replicating in *R. toruloides* to express CRISPR constructs, the most common method for employing CRISPR editing in other fungi (24), requires alternative approaches to express the editing system. One approach to accomplish this would be to directly transform fully assembled Cas9-sgRNA RNP complexes (50). Such an approach has proven successful in the distant basidiomycete relative *Cryptococcus neoformans* (50), suggesting that it may one day prove successful in *R. toruloides.* An alternate approach involving the delivery of DNA coding for CRISPR machinery was explored in this study. We demonstrated that stable genome integration of a Cas9-sgRNA expression cassette using a dominant selectable drug marker is sufficient to achieve gene disruption.

The next major barrier we overcame was the successful expression of sgRNAs intracellularly. This requires a robust RNA Pol-IIII promoter in order to achieve high expression of the guides inside of the nucleus. *R. toruloides,* like many other non-model fungi, have poorly explored Pol-III promoter systems (57). We therefore explored a variety of such promoters, as well as RNA processing elements including ribozymes and self-splicing tRNAs. We found the optimal sgRNA expression system to be tRNA-driven guides, preferably using designs guided from sgRNA prediction programs, such as sgRNA Scorer (38). There is conflicting evidence in the literature as to whether inclusion of a ribozyme element improves sgRNA expression, with some studies finding that it increases editing efficiency (35, 40) while others find the opposite effect (58). Here, our results indicate that ribozyme inclusion is detrimental to Cas9 editing efficiency in *R. toruloides.* Additionally, we demonstrated that multiple functional sgRNAs can be expressed from a single construct using the tRNA processing system described in previous works (44, 45). The low level of editing efficiency in this multiplexed sgRNA design, relative to the editing efficiency levels of the sgRNAs expressed independently (especially *CAR2* sgRNAs), indicates that further optimization of this design could enhance multiplexed gene editing. This could include ensuring high-levels of endogenous expression and efficient processing of the transcript into individual sgRNAs. While the potential for optimization remains, our work towards optimized sgRNA expression provides design guidelines for future CRISPR engineering efforts in *R. toruloides.*

A significant locus-dependent editing efficiency was observed in our study. Despite much of our work focusing on optimization of *URA3* deletion with Cas9, we achieved a relatively low maximum editing efficiency at this locus of 0.62% ± 0.50%. However, deletion of *CAR2* was markedly more successful in even the worst-case scenario (3.4% ± 2.7%), while the best-case scenario resulted in roughly a one-to-one ratio of white to red colonies. The locus-dependency of Cas9 editing efficiency has been noted in other non-yeast eukaryotic systems (23, 59, 60). Combined with the qualitatively large differences observed between editing efficiencies of Cas9 at various *URA3* target locations, optimization of the Cas9 target sequence appears to be particularly important for editing in *R. toruloides.* Transient Cas9 binding events are known to occur much more frequently than actual cleavage occurs, and changes in the protein’s conformation upon binding to the correct target sequence dictate whether DNA cutting actually occurs (61). Furthermore, the high GC content of the *R. toruloides* genome should be taken into consideration (62). A correlation between high GC content and lower Cas9 target specificity has been noted (63), raising the possibility that Cas9 cleavage in *R. toruloides* is (i) more promiscuous or (ii) less effective. A thorough exploration of genome editing efficiencies of Cas9 at various locations in *R. toruloides’* genome would assist in future genome editing endeavors in the organism.

The ability to achieve multiple DNA edits in one round of transformation is an important step forward in *R. toruloides* genome engineering. Thus far, multiple gene edits have been accomplished utilizing multiple rounds of ATMT in which genes are disrupted one at a time; here we have demonstrated that four simultaneous DNA edits can be achieved at once. The sgRNA array could theoretically be expanded to include even more targets. It should be noted that a reduction in editing efficiency may occur with sgRNAs located further downstream in the array. This is supported by the fact that the editing efficiency of *URA3* deletion was relatively similar in the individual and multiplexed targeting constructs (where the targets were located upstream), but the editing efficiency of *CAR2* deletion was substantially lower in the multiplexed targeting construct (where the targets were located downstream) than in the individual targeting case. The multiplexed targeting construct also reveals an interesting phenomenon in which excised DNA between two nearby CRISPR cut sites was inverted and reinserted into the genome. This is potentially due to the relatively high level of NHEJ in *R. toruloides* (15) and indicates that genome integration of exogenous DNA at Cas9 cut-sites may be possible if donor DNA can be supplied concurrently with Cas9 cleavage.

Taken together, our work lays the foundation for Cas9-mediated advanced genome editing in *R. toruloides.* Future efforts could improve upon this framework by employing directed integration of the Cas9 cassette into a specific genetic locus in a Δ*KU70* background or on an ARS-based plasmid to circumvent problems arising from random genome integration in the current strategy. Ultimately, these results should enable rapid engineering of complex *R. toruloides* phenotypes, such as multi-gene pathways to produce biofuels and bioproducts.

## MATERIALS AND METHODS

### Strains and culture conditions

The strain *R. toruloides* IFO0880 (obtained from Biological Resource Center, NITE (NRBC)) was used as the wild type strain for all experiments. Liquid cultures of yeast were grown in YPD (BD Difco) at 30°C and constant 200rpm shaking unless otherwise noted. Solid YPD agar plates were used to grow yeast colonies at 30°C and supplemented with the antibiotics nourseothricin (Werner Bioagents, 100 μg/mL) or 5-fluoroorotic acid (Abcam, 1 mg/mL), or geneticin (VWR, 100 μg/mL) as appropriate.

For all cloning, *Escherichia coli* strains XL1-Blue or DH5α were used to propagate plasmids. *E. coli* were grown in lysogeny broth (LB, BD Difco) at 37°C with 200rpm shaking. Where appropriate, *E*. *coli* media was supplemented with 100 μg/mL ampicillin (or 100 μg/mL carbenicillin in place of ampicillin) or 50 μg/mL kanamycin to maintain plasmids.

### Plasmid construction

The coding sequence of *Streptococcus pyogenes* spCas9 and the SV40 NLS (PKKKRKV) were codon-optimized for expression in *R. toruloides* (GenScript), with a (Gly_)3_ linker included to connect the C-term of spCas9 to the NLS. The fusion protein was placed under expression of the 800bp GAPDH promoter sequence and NOS terminator sequence, which is known to promote strong gene expression in *R. toruloides* (2, 54). The desired optimized sequences were synthesized by GenScript and sequence verified. All plasmids were commercially synthesized excluding plasmids p213 and p227-233, which were constructed in our lab as follows. Plasmid p213 was first constructed by creating two PCR products from p90 using primers 100-103, and subsequently using In-Fusion HD Cloning (Clontech) to stitch the PCR products together. Plasmids p227-233 were created from p213 using primers 104-117 in PCR reactions to create new sgRNA target sequences in individual PCR products, which were subsequently circularized using In-Fusion HD Cloning. A table of strains, plasmids, primers used in this study are in tables S1-3 and are available from the JBEI Registry (https://registry.jbei.org/).

### Transforming DNA preparation

DNA for transformation into *R. toruloides* was prepared from the aforementioned *E. coli* plasmids. To integrate their corresponding Cas9-constructs into the genome, plasmids p90-99 and p184-190 were digested with HindIII, while plasmids pGI103-132 were digested with NdeI. Digestion products were subsequently confirmed using gel electrophoresis and purified using a PCR purification kit (DNA Clean & Concentrator, Zymo Research). These purified products were used directly for transformation. An alternative method was used to integrate the corresponding Cas9-constructs on plasmids p213-233 into the genome. These plasmids were instead PCR amplified using primers 122 and 123 (Table S2), and the PCR products were subsequently confirmed using gel electrophoresis and purified using a PCR purification kit. These purified PCR products were used directly for transformation. Regardless of DNA preparation method, approximately 500 ng of transforming DNA was used for transformation.

### Transformation

Transformation was performed using a modified lithium acetate (LiAc) protocol (14). An individual yeast colony was inoculated into 10 mL YPD medium and grown overnight at 30°C with 200 rpm shaking. The following morning, the OD_600_ of this seed culture was measured and used to inoculate 10mL of fresh YPD to an OD_600_ of 0.2. This culture was grown for another four hours at 30°C with 200 rpm shaking to an OD_600_ of approximately 1.0. Cells were pelleted via centrifugation at 4000g for five minutes, washed twice with 10 mL H_2_O, and washed once with 10 mL 150mM LiAc (Millipore Sigma) at pH 7.6, and resuspended in one mL 150mM LiAc. The pellet was then transferred to 1.5mL microcentrifuge tubes, centrifuged at 8000g for one minute, and the supernatant was removed using a pipette. The wet biomass was then resuspended in 240 μL 50% w/v PEG-4000 (Alfa Aesar), 54 μL 1.0M LiAc, 10 μL of pre-boiled salmon sperm DNA (Invitrogen), and 56 μL of transforming DNA (∼500ng of purified PCR product). The viscous slurry was resuspended via pipetting and incubated at 30°C for 30 minutes, after which 34 μL of 1M dithiothreitol (Millipore Sigma) dissolved in DMSO was added. The transformation was heat shocked at 37°C for 60 minutes, and subsequently pelleted and washed with one mL YPD. The culture was then resuspended in two mL YPD and incubated overnight at 30°C with 200rpm shaking. Cells were pelleted, resuspended in 200 μL YPD, and plated on the appropriate selective media. Utilizing this method to randomly integrate dsDNA via the NHEJ pathway into the *R. toruloides* genome typically provides ∼500 colonies, or a transformation efficiency of ∼1000 transformants/ μg

### Determination of gene edits

*R. toruloides* samples transformed with transforming DNA made from plasmids p90-p99 were plated directly on YPD plates supplemented with 5-FOA and grown for three to four days. Colony forming units were subsequently determined for each transformation. Three transformations were performed for every construct and plated on independent plates to acquire three biological replicates. Three control samples in which no DNA was included in the transformation were also performed to determine the rate of spontaneous 5-FOA resistance.

*R. toruloides* samples transformed with all other transforming DNA (derived from plasmids 184-233 and pGI104-132) harboring a *NAT* selective marker were plated directly on YPD plates supplemented with nourseothricin and grown for two to three days. Three colonies were selected from each transformation and grown overnight. For experiments designed to edit the *URA3* locus, serial dilutions of each culture were plated on both YPD and YPD supplemented with 5-FOA and grown for three to four days. For *CAR2* gene editing experiments, serial dilutions of each culture were plated on YPD. Total colony forming units were determined from serial dilutions providing between 10-1000 countable colonies. Conversion of the red phenotype to white phenotype was used as a proxy for successful editing of the *CAR2* gene to induce a loss of function mutation.

For sequencing, individual colonies were selected from 5-FOA plates and grown overnight in YPD. For Figs. 1, S1 and S2, genomic DNA was prepared using a custom protocol of DNA extraction. For this, 200 μL of yeast culture of approximately 0.40 OD_600_ were centrifuged at maximum speed and the supernatant was removed. Pellets were resuspended in 100 μL of 200mM LiOAc supplemented with 1% SDS and incubated for 15 minutes at 95°C. The samples were supplemented with 300 μL of 96-100% EtOH, vortexed thoroughly, and centrifuged at maximum speed for three minutes. The supernatant was aspirated off, and the pellet was washed once with 70% EtOH. The pellet was resuspended in 100 μL H_2_O and pelleted at maximum speed for 15 seconds. The resulting supernatant containing genomic DNA was recovered for downstream applications. All other genomic DNA was recovered from 100 mg of wet biomass (Quick-DNA Fungal/Bacterial Kit, Zymo Research). Genomic DNA quality was examined by running on an agarose gel, and high-quality DNA was used as a template for PCR. The genomic region around the *URA3* or *CAR2* target sites (see Table S2 for primers) were amplified via PCR and run on an agarose gel. PCR products were purified and submitted for Sanger sequencing (Quintara Biosciences).

## ACKNOWLEDGMENTS

This material is based upon work supported by the U.S. Department of Energy (DOE), Office of Science. Preliminary work in this project establishing experimental protocols was supported by the U.S. DOE, Office of Science, Office of Biological and Environmental Research program under Award Number DE-SC-0012527 to J.M.S. and A.P.A. Work conducted by the DOE Joint BioEnergy Institute was supported by the U.S. DOE, Office of Science, Office of Biological and Environmental Research, through contract DE-AC02-05CH11231. Additional funding was provided by the Toyota Motor Corporation under grant number IQADA62820 to J.M.S. and A.P.A.

The authors declare no competing financial interests related to this work.

P.B.O. wrote the manuscript. M.I., P.B.O., and J.M.S. conceived and designed the experiments. M.I. and J.M.S. designed CRISPR constructs. P.B.O. and M.I. performed experiments and analyzed the data. All authors read, edited, and approved the final manuscript.

## REFERENCES

1. Li Y, Zhao Z (Kent), Bai F. 2007. High-density cultivation of oleaginous yeast Rhodosporidium toruloides Y4 in fed-batch culture. Enzyme Microb Technol 41:312–317. https://10.1016/j.enzmictec.2007.02.008.

2. Yaegashi J, Kirby J, Ito M, Sun J, Dutta T, Mirsiaghi M, Sundstrom ER, Rodriguez A, Baidoo E, Tanjore D, Pray T, Sale K, Singh S, Keasling JD, Simmons BA, Singer SW, Magnuson JK, Arkin AP, Skerker JM, Gladden JM. 2017. Rhodosporidium toruloides: A new platform organism for conversion of lignocellulose into terpene biofuels and bioproducts. Biotechnol Biofuels 10:1–13. https://10.1186/s13068-017-0927-5.

3. Xu J, Liu D. 2017. Exploitation of genus Rhodosporidium for microbial lipid production. World J Microbiol Biotechnol 33:1–13. https://10.1007/s11274-017-2225-6.

4. Chaturvedi S, Bhattacharya A, Khare SK. 2018. Trends in oil production from oleaginous yeast using biomass: biotechnological potential and constraints. Appl Biochem Microbiol 54:361–369. https://10.1134/S000368381804004X.

5. Singh G, Jawed A, Paul D, Bandyopadhyay KK, Kumari A, Haque S. 2016. Concomitant production of lipids and carotenoids in Rhodosporidium toruloides under osmotic stress using response surface methodology. Front Microbiol 7:1–13. https://10.3389/fmicb.2016.01686.

6. Hu C, Zhao X, Zhao J, Wu S, Zhao ZK. 2009. Effects of biomass hydrolysis by-products on oleaginous yeast Rhodosporidium toruloides. Bioresour Technol 100:4843–4847. https://10.1016/j.biortech.2009.04.041.

7. Huang Q, Wang Q, Gong Z, Jin G, Shen H, Xiao S, Xie H, Ye S, Wang J, Zhao ZK. 2013. Effects of selected ionic liquids on lipid production by the oleaginous yeast Rhodosporidium toruloides. Bioresour Technol 130:339–344. https://10.1016/j.biortech.2012.12.022.

8. Sundstrom E, Yaegashi J, Yan J, Masson F, Papa G, Rodriguez A, Mirsiaghi M, Liang L, He Q, Tanjore D, Pray TR, Singh S, Simmons B, Sun N, Magnuson J, Gladden J. 2018. Demonstrating a separation-free process coupling ionic liquid pretreatment, saccharification, and fermentation with: Rhodosporidium toruloides to produce advanced biofuels. Green Chem 20:2870–2879. https://10.1039/c8gc00518d.

9. Díaz T, Fillet S, Campoy S, Vázquez R, Viña J, Murillo J, Adrio JL. 2018. Combining evolutionary and metabolic engineering in Rhodosporidium toruloides for lipid production with non-detoxified wheat straw hydrolysates. Appl Microbiol Biotechnol 102:3287–3300. https://10.1007/s00253-018-8810-2.

10. Marella ER, Holkenbrink C, Siewers V, Borodina I. 2018. Engineering microbial fatty acid metabolism for biofuels and biochemicals. Curr Opin Biotechnol 50:39–46. https://10.1016/j.copbio.2017.10.002.

11. Park Y, Nicaud J, Ledesma-amaro R. 2018. The engineering potential of Rhodosporidium toruloides as a workhorse for biotechnological applications. Trends Biotechnol 36:304–317. https://10.1016/j.tibtech.2017.10.013.

12. Ko JK, Lee SM. 2018. Advances in cellulosic conversion to fuels: engineering yeasts for cellulosic bioethanol and biodiesel production. Curr Opin Biotechnol 50:72–80. https://10.1016/j.copbio.2017.11.007.

13. Shi S, Zhao H. 2017. Metabolic engineering of oleaginous yeasts for production of fuels and chemicals. Front Microbiol 8:1–16. https://10.3389/fmicb.2017.02185.

14. Tsai YY, Ohashi T, Kanazawa T, Polburee P, Misaki R, Limtong S, Fujiyama K. 2017. Development of a sufficient and effective procedure for transformation of an oleaginous yeast, Rhodosporidium toruloides DMKU3-TK16. Curr Genet 63:359–371. https://10.1007/s00294-016-0629-8.

15. Liu H, Jiao X, Wang Y, Yang X, Sun W, Wang J, Zhang S, Zhao ZK. 2017. Fast and efficient genetic transformation of oleaginous yeast Rhodosporidium toruloides by using electroporation. FEMS Yeast Res 17:1–11. https://10.1093/femsyr/fox017.

16. Lin X, Wang Y, Zhang S, Zhu Z, Zhou YJ, Yang F, Sun W, Wang X, Zhao ZK. 2014. Functional integration of multiple genes into the genome of the oleaginous yeast Rhodosporidium toruloides. FEMS Yeast Res 14:547–555. https://10.1111/1567-1364.12140.

17. Coradetti ST, Pinel D, Geiselman GM, Ito M, Mondo SJ, Reilly MC, Cheng Y-F, Bauer S, Grigoriev I V, Gladden JM, Simmons BA, Brem RB, Arkin AP, Skerker JM. 2018. Functional genomics of lipid metabolism in the oleaginous yeast Rhodosporidium toruloides. Elife 7:e32110. https://10.7554/eLife.32110.

18. Koh CMJ, Liu Y, Moehninsi, Du M, Ji L. 2014. Molecular characterization of KU70 and KU80 homologues and exploitation of a KU70-deficient mutant for improving gene deletion frequency in Rhodosporidium toruloides. BMC Microbiol 14:53–68. https://10.1186/1471-2180-14-50.

19. Zhang S, Ito M, Skerker JM, Arkin AP, Rao C V. 2016. Metabolic engineering of the oleaginous yeast Rhodosporidium toruloides IFO0880 for lipid overproduction during high-density fermentation. Appl Microbiol Biotechnol 100:9393–9405. https://10.1007/s00253-016-7815-y.

20. Hsu PD, Lander ES, Zhang F. 2014. Development and applications of CRISPR-Cas9 for genome engineering. Cell 157:1262–1278. https://10.1016/j.cell.2014.05.010.

21. Ledford H. 2015. CRISPR, the disruptor. Nature 522:20–24. https://10.1038/522020a.

22. Jinek M, Chylinski K, Fonfara I, Hauer M, Doudna JA, Charpentier E. 2012. A programmable dual-RNA-guided DNA endonuclease in adaptive bacterial immunity. Science 337:816–821. https://10.1126/science.1225829.

23. Langner T, Kamoun S, Belhaj K. 2018. CRISPR crops: plant genome editing toward disease resistance. Annu Rev Phytopathol 56:479–512. https://10.1146/annurev-phyto-080417-050158.

24. Shi TQ, Liu GN, Ji RY, Shi K, Song P, Ren LJ, Huang H, Ji XJ. 2017. CRISPR/Cas9-based genome editing of the filamentous fungi: the state of the art. Appl Microbiol Biotechnol 101:7435–7443. https://10.1007/s00253-017-8497-9.

25. Deng H, Gao R, Liao X, Cai Y. 2017. CRISPR system in filamentous fungi: Current achievements and future directions. Gene 627:212–221. https://10.1016/j.gene.2017.06.019.

26. Blazeck J, Hill A, Liu L, Knight R, Miller J, Pan A, Otoupal P, Alper HS. 2014. Harnessing Yarrowia lipolytica lipogenesis to create a platform for lipid and biofuel production. Nat Commun 5:3131. https://10.1038/ncomms4131.

27. Adrio JL. 2017. Oleaginous yeasts: Promising platforms for the production of oleochemicals and biofuels. Biotechnol Bioeng 114:1915–1920. https://10.1002/bit.26337.

28. Liu L, Otoupal P, Pan A, Alper HS. 2014. Increasing expression level and copy number of a Yarrowia lipolytica plasmid through regulated centromere function. FEMS Yeast Res https://10.1111/1567-1364.12201.

29. Markham KA, Alper HS. 2018. Synthetic Biology Expands the Industrial Potential of Yarrowia lipolytica. Trends Biotechnol 1–11. https://10.1016/j.tibtech.2018.05.004.

30. Zetsche B, Heidenreich M, Mohanraju P, Fedorova I, Kneppers J, Degennaro EM, Winblad N, Choudhury SR, Abudayyeh OO, Gootenberg JS, Wu WY, Scott DA, Severinov K, Van Der Oost J, Zhang F. 2017. Multiplex gene editing by CRISPR-Cpf1 using a single crRNA array. Nat Biotechnol 35:31–34. https://10.1038/nbt.3737.

31. Verwaal R, Buiting-Wiessenhaan N, Dalhuijsen S, Roubos JA. 2018. CRISPR/Cpf1 enables fast and simple genome editing of Saccharomyces cerevisiae. YEAST 35:201–211. https://10.1002/yea.3278.

32. Krappmann S. 2007. Gene targeting in filamentous fungi: the benefits of impaired repair. Fungal Biol Rev 21:25–29. https://10.1016/j.fbr.2007.02.004.

33. Dicarlo JE, Norville JE, Mali P, Rios X, Aach J, Church GM. 2013. Genome engineering in Saccharomyces cerevisiae using CRISPR-Cas systems. Nucleic Acids Res 41:4336–4343. https://10.1093/nar/gkt135.

34. Krappmann S. 2017. CRISPR-Cas9, the new kid on the block of fungal molecular biology. Med Mycol 55:16–23. https://10.1093/mmy/myw097.

35. Ryan OW, Skerker JM, Maurer MJ, Li X, Tsai JC, Poddar S, Lee ME, DeLoache W, Dueber JE, Arkin AP, Cate JHD. 2014. Selection of chromosomal DNA libraries using a multiplex CRISPR system. Elife 3:1–15. https://10.7554/eLife.03703.

36. Alani E, Cao L, Kleckner N. 1987. A method for gene disruption that allows repeated use of URA3 selection in the construction of multiply disrupted yeast strains. Genetics 116:541–545. https://10.1534/genetics.112.541.test.

37. Radzisheuskaya A, Shlyueva D, Müller I, Helin K. 2016. Optimizing sgRNA position markedly improves the efficiency of CRISPR/dCas9-mediated transcriptional repression. Nucleic Acids Res 44. https://10.1093/nar/gkw583.

38. Chari R, Yeo NC, Chavez A, Church GM. 2017. SgRNA Scorer 2.0: a species-independent model to predict CRISPR/Cas9 activity. ACS Synth Biol 6:902–904. https://10.1021/acssynbio.6b00343.

39. Braman JC. 2018. Synthetic Biology Methods and ProtocolsMethods in Molecular Biology.

40. de Vries ARG, de Groot PA, van den Broek M, Daran JMG. 2017. CRISPR-Cas9 mediated gene deletions in lager yeast Saccharomyces pastorianus. Microb Cell Fact 16:1–18. https://10.1186/s12934-017-0835-1.

41. Schwartz CM, Hussain MS, Blenner M, Wheeldon I. 2016. Synthetic RNA Polymerase III Promoters Facilitate High-Efficiency CRISPR-Cas9-Mediated Genome Editing in Yarrowia lipolytica. ACS Synth Biol 5:356–359. https://10.1021/acssynbio.5b00162.

42. Song L, Ouedraogo JP, Kolbusz M, Nguyen TTM, Tsang A. 2018. Efficient genome editing using tRNA promoter-driven CRISPR/Cas9 gRNA in Aspergillus Niger. PLoS One 13:1–17. https://10.1371/journal.pone.0202868.

43. Landolfo S, Ianiri G, Camiolo S, Porceddu A, Mulas G, Chessa R, Zara G, Mannazzu I. 2018. CAR gene cluster and transcript levels of carotenogenic genes in Rhodotorula mucilaginosa. Microbiol (United Kingdom) 164:78–87. https://10.1099/mic.0.000588.

44. Xie K, Minkenberg B, Yang Y. 2015. Boosting CRISPR/Cas9 multiplex editing capability with the endogenous tRNA-processing system. Proc Natl Acad Sci 112:3570–3575. https://10.1073/pnas.1420294112.

45. Nødvig CS, Hoof JB, Kogle ME, Jarczynska ZD, Lehmbeck J, Klitgaard DK, Mortensen UH. 2018. Efficient oligo nucleotide mediated CRISPR-Cas9 gene editing in Aspergilli. Fungal Genet Biol 115:78–89. https://10.1016/j.fgb.2018.01.004.

46. Wang H, Yang H, Shivalila CS, Dawlaty MM, Cheng AW, Zhang F, Jaenisch R. 2013. One-step generation of mice carrying mutations in multiple genes by CRISPR/cas-mediated genome engineering. Cell 153:910–918. https://10.1016/j.cell.2013.04.025.

47. Liu Q, Gao R, Li J, Lin L, Zhao J, Sun W, Tian C. 2017. Development of a genome-editing CRISPR/Cas9 system in thermophilic fungal Myceliophthora species and its application to hyper-cellulase production strain engineering. Biotechnol Biofuels 10:1–14. https://10.1186/s13068-016-0693-9.

48. Ran FA, Hsu PD, Wright J, Agarwala V, Scott DA, Zhang F. 2013. Genome engineering using the CRISPR-Cas9 system. Nat Protoc 8:2281–2308. https://10.1038/nprot.2013.143.

49. Sarkari P, Marx H, Blumhoff ML, Mattanovich D, Sauer M, Steiger MG. 2017. An efficient tool for metabolic pathway construction and gene integration for Aspergillus niger. Bioresour Technol 245:1327–1333. https://10.1016/j.biortech.2017.05.004.

50. Wang P. 2018. Two distinct approaches for CRISPR-Cas9-mediated gene editing in Cryptococcus neoformans and related species. mSphere 3:e00208–18. https://10.1128/mSphereDirect.00208-18.

51. Nagy G, Szebenyi C, Csernetics Á, Vaz AG, Tóth EJ, Vágvölgyi C, Papp T. 2017. Development of a plasmid free CRISPR-Cas9 system for the genetic modification of Mucor circinelloides. Sci Rep 7:1–10. https://10.1038/s41598-017-17118-2.

52. Sun W, Yang X, Wang X, Jiao X, Zhang S, Luan Y, Zhao ZK. 2018. Developing a flippase-mediated maker recycling protocol for the oleaginous yeast Rhodosporidium toruloides. Biotechnol Lett 40:933–940. https://10.1007/s10529-018-2542-3.

53. Johns AMB, Love J, Aves SJ. 2016. Four inducible promoters for controlled gene expression in the oleaginous yeast Rhodotorula toruloides. Front Microbiol 7:1585. https://10.3389/fmicb.2016.01666.

54. Liu Y, Koh CMJ, Sun L, Hlaing MM, Du M, Peng N, Ji L. 2013. Characterization of glyceraldehyde-3-phosphate dehydrogenase gene RtGPD1 and development of genetic transformation method by dominant selection in oleaginous yeast Rhodosporidium toruloides. Appl Microbiol Biotechnol 97:719–729. https://10.1007/s00253-012-4223-9.

55. Wang Y, Lin X, Zhang S, Sun W, Ma S, Zhao ZK. 2016. Cloning and evaluation of different constitutive promoters in the oleaginous yeast Rhodosporidium toruloides. Yeast 33:99–106. https://10.1002/yea.3145.

56. Liu Y, Yap SA, Koh CMJ, Ji L. 2016. Developing a set of strong intronic promoters for robust metabolic engineering in oleaginous Rhodotorula (Rhodosporidium) yeast species. Microb Cell Fact 15:1–9. https://10.1186/s12934-016-0600-x.

57. Morse NJ, Wagner JM, Reed KB, Gopal MR, Lauffer LH, Alper HS. 2018. T7 polymerase expression of guide RNAs in vivo allows exportable CRISPR-Cas9 editing in multiple yeast hosts. ACS Synth Biol 7:1075–1084. https://10.1021/acssynbio.7b00461.

58. Apel AR, D’Espaux L, Wehrs M, Sachs D, Li RA, Tong GJ, Garber M, Nnadi O, Zhuang W, Hillson NJ, Keasling JD, Mukhopadhyay A. 2017. A Cas9-based toolkit to program gene expression in Saccharomyces cerevisiae. Nucleic Acids Res 45:496–508. https://10.1093/nar/gkw1023.

59. Hsu PD, Scott D a, Weinstein J a, Ran FA, Konermann S, Agarwala V, Li Y, Fine EJ, Wu X, Shalem O, Cradick TJ, Marraffini L a, Bao G, Zhang F. 2013. DNA targeting specificity of RNA-guided Cas9 nucleases. Nat Biotechnol 31:827–32. https://10.1038/nbt.2647.

60. Merkle FT, Neuhausser WM, Santos D, Valen E, Gagnon JA, Maas K, Sandoe J, Schier AF, Eggan K. 2015. Efficient CRISPR-Cas9-mediated generation of knockin human pluripotent stem cells lacking undesired mutations at the targeted locus. Cell Rep 11:875–883. https://10.1016/j.celrep.2015.04.007.

61. Sternberg SH, Lafrance B, Kaplan M, Doudna JA. 2015. Conformational control of DNA target cleavage by CRISPR-Cas9. Nature 527:110–113. https://10.1038/nature15544.

62. Sambles C, Middelhaufe S, Soanes D, Kolak D, Lux T, Moore K, Matoušková P, Parker D, Lee R, Love J, Aves SJ. 2017. Genome sequence of the oleaginous yeast Rhodotorula toruloides strain CGMCC 2.1609. Genomics Data 13:1–2. https://10.1016/j.gdata.2017.05.009.

63. Tsai SQ, Zheng Z, Nguyen NT, Liebers M, Topkar V V., Thapar V, Wyvekens N, Khayter C, Iafrate AJ, Le LP, Aryee MJ, Joung JK. 2015. GUIDE-seq enables genome-wide profiling of off-target cleavage by CRISPR-Cas nucleases. Nat Biotechnol 33:187–198. https://10.1038/nbt.3117.

