## Supplementary Tables and Figures for "Multiplexed CRISPR-Cas9 based genome editing of *Rhodosporidium toruloides*"

**Table S1** Plasmids used in this study

| Name | Description | Source |
|---|---|---|
| 90 | Cas9-sgRNA(URA3-1,SC-SNR52pr,Ribozyme) | Genscript |
| 91 | Cas9-sgRNA(URA3-1,SC-tRNA$^{Phe}$,Ribozyme) | Genscript |
| 92 | Cas9-sgRNA(URA3-1,RT-tRNA$^{Phe}$,Ribozyme) | Genscript |
| 93 | Cas9-sgRNA(URA3-1,RT-tRNA$^{Pro}$,Ribozyme) | Genscript |
| 94 | Cas9-sgRNA(URA3-1,RT-tRNA$^{Tyr}$,Ribozyme) | Genscript |
| 95 | Cas9-sgRNA(URA3-2,SC-SNR52pr,Ribozyme) | Genscript |
| 96 | Cas9-sgRNA(URA3-2,SC-tRNA$^{Phe}$,Ribozyme) | Genscript |
| 97 | Cas9-sgRNA(URA3-2,RT-tRNA$^{Phe}$,Ribozyme) | Genscript |
| 98 | Cas9-sgRNA(URA3-2,RT-tRNA$^{Pro}$,Ribozyme) | Genscript |
| 99 | Cas9-sgRNA(URA3-2,tRNA$^{Tyr}$,Ribozyme) | Genscript |
| 184 | Cas9-sgRNA(URA3-1,SC-SNR52pr,Ribozyme)-NAT | Genscript |
| 185 | Cas9-sgRNA(URA3-1,SC-tRNA$^{Phe}$,Ribozyme)-NAT | Genscript |
| 186 | Cas9-sgRNA(URA3-1,RT-tRNA$^{Phe}$,Ribozyme)-NAT | Genscript |
| 187 | Cas9-sgRNA(URA3-1,RT-tRNA$^{Pro}$,Ribozyme)-NAT | Genscript |
| 188 | Cas9-sgRNA(URA3-1,RT-tRNA$^{Tyr}$,Ribozyme)-NAT | Genscript |
| 190 | Cas9-eGFP-sgRNA(URA3-1,SC-SNR52pr,Ribozyme)-NAT | Genscript |
| 213 | Cas9-sgRNA(URA3-1,SC-SNR52pr,Ribozyme)-NAT | This Study |
| 227 | Cas9-sgRNA(URA3-3,SC-SNR52pr,Ribozyme)-NAT | This Study |
| 228 | Cas9-sgRNA(URA3-4,SC-SNR52pr,Ribozyme)-NAT | This Study |
| 229 | Cas9-sgRNA(URA3-5,SC-SNR52pr,Ribozyme)-NAT | This Study |
| 230 | Cas9-sgRNA(URA3-6,SC-SNR52pr,Ribozyme)-NAT | This Study |
| 231 | Cas9-sgRNA(URA3-7,SC-SNR52pr,Ribozyme)-NAT | This Study |
| 232 | Cas9-sgRNA(URA3-8,SC-SNR52pr,Ribozyme)-NAT | This Study |
| 233 | Cas9-sgRNA(URA3-9,SC-SNR52pr,Ribozyme)-NAT | This Study |
| pGI104 | Cas9-sgRNA(URA3-1,SC-SNR52pr,YL-tRNA$^{Gly}$)-NAT | Genscript |
| pGI105 | Cas9-sgRNA(URA3-1,YL-SNR52pr)-NAT | Genscript |
| pGI106 | Cas9-sgRNA(URA3-1,YL-tRNA$^{Gly}$)-NAT | Genscript |
| pGI107 | Cas9-sgRNA(URA3-1,SC-tRNA$^{Phe}$)-NAT | Genscript |
| pGI108 | Cas9-sgRNA(URA3-1,SC-SNR52pr)-NAT | Genscript |
| pGI110 | Cas9-sgRNA(URA3-1,RT-tRNA$^{Tyr}$)-NAT | Genscript |
| pGI119 | Cas9-sgRNA(URA3-4,RT-tRNA$^{Tyr}$)-NAT | Genscript |
| pGI120 | Cas9-sgRNA(CAR2-1,RT-tRNA$^{Tyr}$)-NAT | Genscript |
| pGI121 | Cas9-sgRNA(CAR2-2,RT-tRNA$^{Tyr}$)-NAT | Genscript |
| pGI122 | Cas9-sgRNA(CAR2-3,RT-tRNA$^{Tyr}$)-NAT | Genscript |
| pGI123 | Cas9-sgRNA(CAR2-4,RT-tRNA$^{Tyr}$)-NAT | Genscript |
| pGI132 | Cas9-sgRNA(URA3-3+4,Car2-1+3,RT-tRNA$^{Tyr}$)-NAT | Genscript |

**Table S2** Primers used in this study

| Name | Description | Sequence (5' – 3') |
|---|---|---|
| 100 | Constructing p213 Fwd Fragment 1 | acctcgctcgcgagaCGTTGCTGGCGTTTTTCCATAG |
| 101 | Constructing p213 Rev Fragment 1 | TTATCTTTTCAAAGAAAGGTGGCACTTTTCGGGGAAATG |
| 102 | Constructing p213 Fwd Fragment 2 | AAAGTGCCACCTTTCTTTGAAAAGATAATGTATGATTATGCTTTCACTC |
| 103 | Constructing p213 Rev Fragment 2 | AAAACGCCAGCAACGtctcgcgagcgaggttggct |
| 104 | Constructing p227 Fwd | CTCGTCACGTCAACCAAAGTCCCATTCGCCACCC |
| 105 | Constructing p227 Rev | ACGTGACGAGCAAGGGTTTTAGAGCTAGAAATAGCAAGTTAAAATAAGGCTAGTC |
| 106 | Constructing p228 Fwd | TCCCCGGCGAGGGTAAAAGTCCCATTCGCCACCC |
| 107 | Constructing p228 Rev | TCGCCGGGGACCAGGGTTTTAGAGCTAGAAATAGCAAGTTAAAATAAGGCTAGTC |
| 108 | Constructing p229 Fwd | TGCTGGACGAGATCCAAAGTCCCATTCGCCACCC |
| 109 | Constructing p229 Rev | TCGTCCAGCAACTGCGTTTTAGAGCTAGAAATAGCAAGTTAAAATAAGGCTAGTC |
| 110 | Constructing p230 Fwd | TCGCCCTTCGAGTCGAAAGTCCCATTCGCCACCC |
| 111 | Constructing p230 Rev | CGAAGGGCGACGGCAGTTTTAGAGCTAGAAATAGCAAGTTAAAATAAGGCTAGTC |
| 112 | Constructing p231 Fwd | AACACCGCCAGCCGTAAAGTCCCATTCGCCACCC |
| 113 | Constructing p231 Rev | TGGCGGTGTTGCGCCGTTTTAGAGCTAGAAATAGCAAGTTAAAATAAGGCTAGTC |
| 114 | Constructing p232 Fwd | CCTCTGCGATCCTGAAAAGTCCCATTCGCCACCC |
| 115 | Constructing p232 Rev | ATCGCAGAGGCTGCTGTTTTAGAGCTAGAAATAGCAAGTTAAAATAAGGCTAGTC |
| 116 | Constructing p233 Fwd | AGAAATCGTGCTTGTAAAGTCCCATTCGCCACCC |
| 117 | Constructing p233 Rev | CACGATTTCTTGATCGTTTTAGAGCTAGAAATAGCAAGTTAAAATAAGGCTAGTC |
| 118 | Car2 Genomic DNA PCR Fwd | CCACCTCCGCTGGACTATCC |
| 119 | Car2 Genomic DNA PCR Rev | GGCAACTCGGCGGAGATACTC |
| 120 | Ura3 Genomic DNA PCR Fwd | CCGTCTCTTCCCGCTCATC |
| 121 | Ura3 Genomic DNA PCR Rev | CAGCTCCGTCCTTCGAGATG |
| 122 | Transforming DNA PCR Fwd | ACTATCGTCTTGAGTCCAACCCGGTAAGAC |
| 123 | Transforming DNA PCR Rev | TGAATCGACAAGATCTCGATAGCCGCCTGCAAGTCCTCGATCTCGCGAGCGAGGTTGG CT |

**Table S3** sgRNA guide target sequences

| Description | Sequence (5' – 3') |
|---|---|
| Ura3 Target1 | CCAAGAACGTCGAGGCGGCG |
| Ura3 Target2 | GAAGACCAACCTCTGCGTCT |
| Ura3 Target3 | GGTTGACGTGACGAGCAAGG |
| Ura3 Target4 | TACCCTCGCCGGGGACCAGG |
| Ura3 Target5 | GGATCTCGTCCAGCAACTGC |
| Ura3 Target6 | CGACTCGAAGGGCGACGGCA |
| Ura3 Target7 | ACGGCTGGCGGTGTTGCGCC |
| Ura3 Target8 | TCAGGATCGCAGAGGCTGCT |
| Ura3 Target9 | ACAAGCACGATTTCTTGATC |
| Car1 Target1 | CAGATCGAGGCGGCCTTGCC |
| Car2 Target2 | CGCACAGCGACAACGCATCG |
| Car3 Target3 | CCTTCAGAAAGCTTAGGACA |
| Car4 Target4 | GGACGGCGTGGACGACTGGG |

### Supplementary Figures

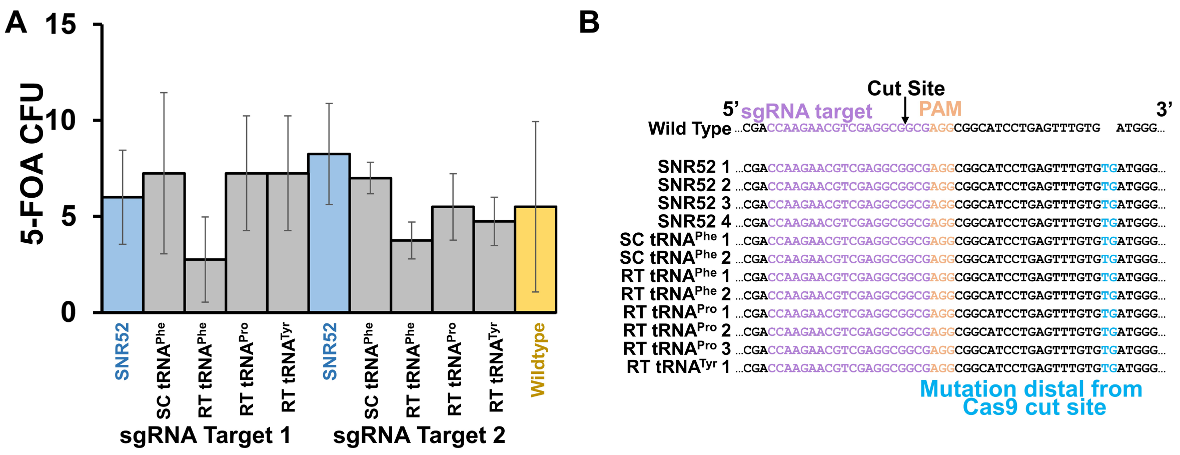

**Figure S1.** (A) Colony forming units (CFUs) of *R. toruloides* upon transformation of various CRISPR expression constructs in which the sgRNA is expressed under various promoters, using the approach outlined in Figure 1A. The original promoter is highlighted in blue, while transformation with no transforming DNA in wildtype *R. toruloides* is highlighted in yellow. The source of the promoter sequence as either *S. cerevisiae* (Sc) or *R. toruloides* (Rt) are listed in parentheses. Error bars represent the standard deviation of biological triplicates. A two-tailed type II student's t-test was performed comparing the CFU of wildtype to the other ten samples, but no significant differences were observed. (B) Sequencing results of various replicates selected from transformed plates of sgRNA Target 1. The Cas9 cut site is indicated, while INDELs are highlighted in blue.

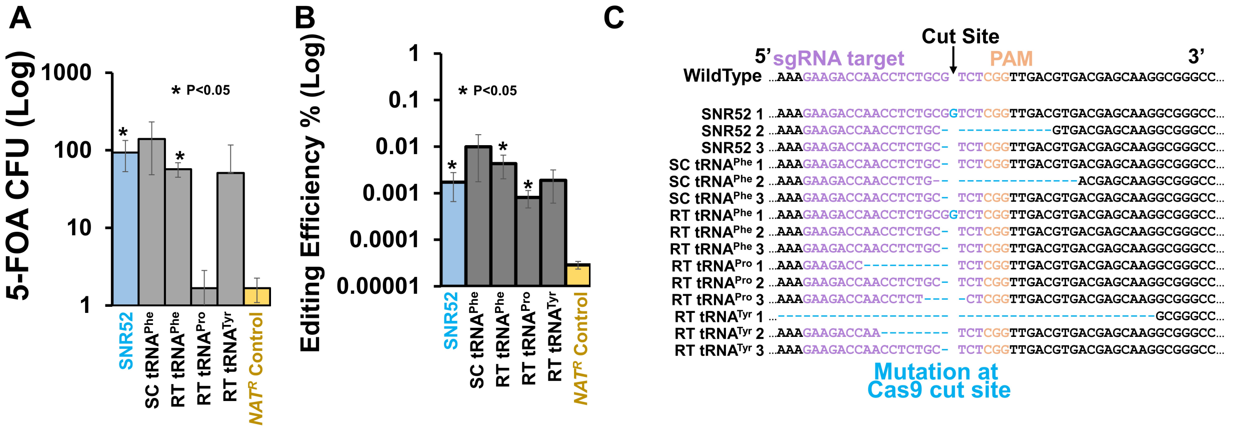

**Figure S2.** (A) Colony forming units (CFUs) of *R. toruloides* upon transformation of various CRISPR expression constructs in which the sgRNA is expressed under various promoters, using the approach outlined in Figure 1D using sgRNA Target 2. The original promoter is highlighted in blue, while transformation with a $NAT^R$ cassette excluding the Cas9 and sgRNA sequences into wildtype *R. toruloides* is highlighted in yellow. The source of the promoter sequence as either *S. cerevisiae* (SC) or *R. toruloides* (RT) are listed in parentheses. (B) Editing efficiencies of each construct, determined as the ratio of 5-FOA CFU to NAT CFU. The low level of construct Pro tRNA (RT) 5-FOA CFU was due to a low number of overall cells used in the transformation; when this was taken into account, editing efficiencies of this construct were similar to the original construct. Additionally, the "editing efficiency" due to spontaneous mutation of 5-FOA resistance was calculated from the $NAT^R$ control transformation. Error bars represent the standard deviation of biological triplicates. P-values were calculated using a two-tailed type II student's t-test with comparison to the $NAT^R$ control. (C) Sequencing results of various replicates selected from transformed plates. The Cas9 cut site is indicated, while INDELs are highlighted in blue.

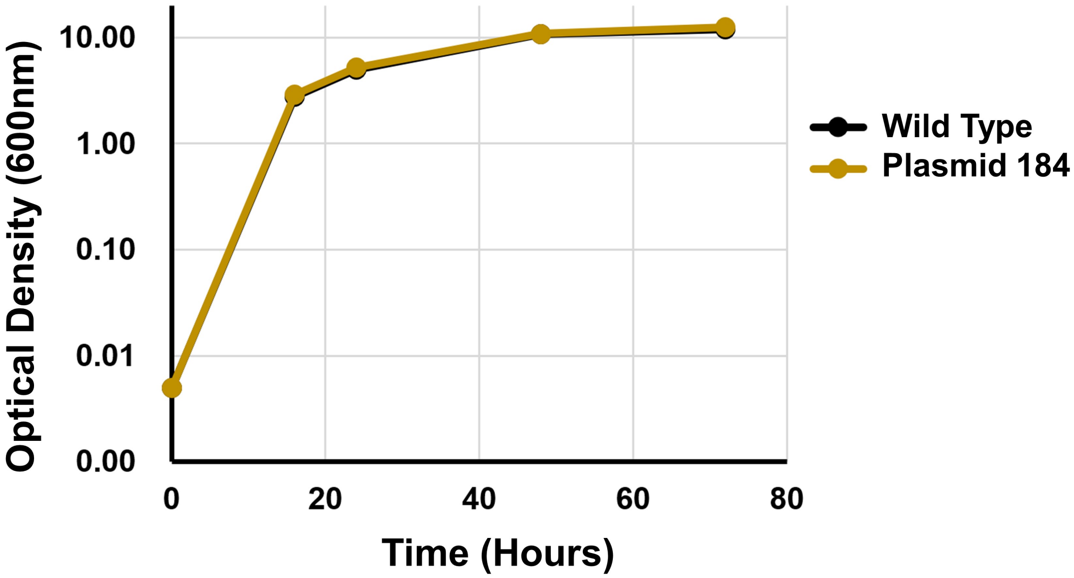

**Figure S3.** Growth of wildtype *R. toruloides* in comparison to *R. toruloides* harboring a genome-integrated Cas9 expression cassette. Growth was performed in YPD supplemented with no antibiotics.

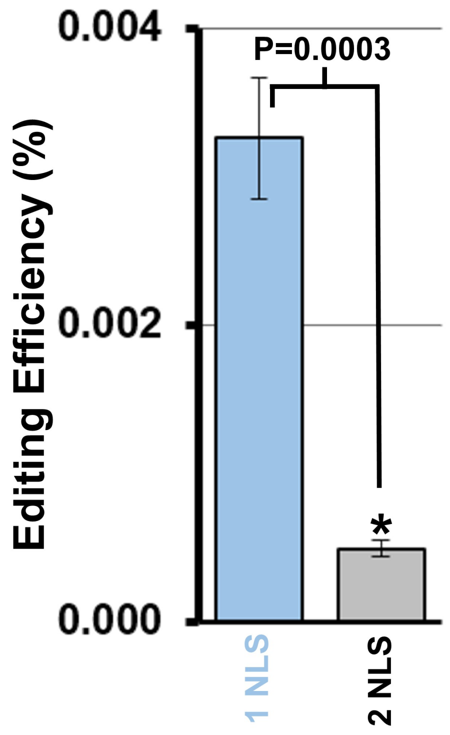

**Figure S4. Effect of additional NLS on Cas9 editing efficiency.** We first used the SV40 NLS as it has been shown to work well in a variety of fungal systems. However, we thought editing efficiency might be improved by adding an additional NLS to the Cas9 sequence. To test this, we included a second SV40 NLS downstream of the first. However, this change significantly ($P = 0.0003$) reduced editing efficiency roughly ~7 fold, from 0.0032 ± 0.0004% to 0.00049 ± 0.00006%. We therefore do not recommend this approach in *R. toruloides*.
